# PP2A activation drives aberrant macropinocytosis and cell death in pancreatic ductal adenocarcinoma

**DOI:** 10.1101/2025.07.14.664742

**Authors:** Garima Baral, Claire M. Pfeffer, Sara N. Filippelli, Indiraa Doraivel, Brittany N. Heil, Lauren E. Gartenhaus, Kasi Hansen, Joy Wu, Goutham Narla, Emma H. Doud, Amber L. Mosley, Shalini T. Low-Nam, Brittany L. Allen-Petersen

## Abstract

Pancreatic cancer is highly aggressive with a five-year survival rate of just 13%. Metabolic rewiring in response to oncogenic signals plays a critical role in pancreatic ductal adenocarcinoma (PDAC) survival, tumor growth, and metastasis. These alterations make PDAC tumors dependent on anabolic metabolism for survival, highlighting a unique vulnerability that can be therapeutically exploited. However, during nutrient deprivation, PDAC cells can circumvent this vulnerability by engulfing extracellular fluids to replenish amino acids in a process called, macropinocytosis. This process can be induced downstream of oncogenic KRAS expression, a small GTPase that is almost universally mutated in PDAC patients. The inhibition of macropinocytosis *in vivo* reduces PDAC tumor growth, emphasizing the importance of this pathway to cancer cell survival. However, the signaling mechanisms that regulate this process remain poorly understood. Protein phosphatase 2A (PP2A) is a heterotrimeric complex that regulates a wide variety of cell signaling pathways, including KRAS, and is commonly dysregulated in human PDAC tumors. Here, we show that acute PP2A activation prevents macropinosome processing leading to cell death. Furthermore, we demonstrate that PP2A posttranslationally regulates the lipid kinase, PIKfyve, a key regulator of macropinosome-lysosome fusion. Finally, we determine that PP2A activating compounds can function synergistically with metabolic inhibitors, supporting a new therapeutic strategy in this aggressive and deadly cancer. Together, our results implicate PP2A as a critical suppressor of PDAC metabolic plasticity and highlight the use of PP2A activating compounds to prevent PDAC nutrient scavenging.

## INTRODUCTION

The majority of pancreatic ductal adenocarcinoma (PDAC) patients present with late-stage, metastatic disease that is often resistant to therapeutics, resulting in the lowest 5-year survival rate of all major cancers^1,2^. While significant advances have been made in the development of therapeutic strategies, most PDAC patients remain refractory to standard of care chemotherapy^3^. One of the major contributing factors to therapeutic resistance in PDAC is the dense fibrotic stroma that accounts for up to 90% of the total tumor mass^2,4,5^. This aggressive tumor microenvironment causes poor vascularization, increased interstitial fluid pressure, and reduced nutrient availability^6,7^. The limited supply of essential nutrients in combination with high metabolic demand results in PDAC cells employing alternative nutrient scavenging strategies like autophagy and macropinocytosis for survival^8–12^. During macropinocytosis, PDAC cells take up bulk nutrients from the microenvironment through the non-selective internalization of extracellular fluid. The increase in nutrient availability through macropinocytosis is known to mediate tumor cell survival and increase PDAC tumor growth, underscoring the importance of this metabolic pathway^10^. Therefore, targeting this scavenging mechanism provides unique therapeutic opportunities for PDAC patients.

Macropinocytosis is mediated in part by the small GTPase, Kirsten rat sarcoma virus (*KRAS*)^9,10,12^. *KRAS* mutations are considered to be major drivers of this aggressive disease and promote metabolic rewiring, proliferation, and survival^13–15^. During macropinocytosis, oncogenic KRAS expression is known to activate Rac family small GTPase 1 (RAC1), facilitating actin polymerization and the formation of membrane ruffles^9,16^. Extracellular proteins are then encapsulated and internalized into vesicles called macropinosomes. Once inside the cell, macropinosomes undergo a maturation process that culminates in the fusion of macropinosomes with lysosomes, where bulk nutrients are digested and recycled. While macropinosome maturation and fusion with the lysosome are known to be regulated by changes in the posttranslational modifications of membrane lipids^17^, the contribution of protein phosphorylation to the fate of macropinosomes remains understudied.

Protein Phosphatase 2A (PP2A) is a heterotrimeric protein that accounts for a substantial portion of serine/threonine phosphatase activity in cells^18^. The PP2A holoenzyme is composed of a scaffolding A subunit, a regulatory B subunit, and a catalytic C subunit. The PP2A B subunits fall into four major families (B (B55), B’ (B56), B’’ and B’’’) and the distinct B subunit bound to PP2A A/C dimer determines substrate specificity of the PP2A holoenzyme^19,20^. The indirect activation of PP2A by the synthetic sphingolipid, SH-BC-893 has been shown to promote macropinocytosis^21^. Yet, the specific mechanism by which PP2A regulates the fate of macropinosomes, and the regulatory subunits involved in this regulation remains unclear.

Here, we show that activation of PP2A alters macropinosome maturation in PDAC cells, primarily through the regulatory subunit B56α. In contrast to the known roles for macropinocytosis in cell survival, pharmacological activation of PP2A disrupts macropinosome-lysosome fusion, leading to macropinosome accumulation and cell death. Furthermore, we demonstrate that the PP2A regulatory subunit, B56α, directly associates with a protein involved in the macropinosome maturation process. Given the negative regulation of the macropinocytosis cascade by PP2A, we explored the impact of pharmacological PP2A activation in the combinatorial targeting of nutrient acquisition pathways. In particular, we determine that pharmacological PP2A activation functions synergistically with the amino acid transporter inhibitor, V-9302, *in vitro* and significantly decreases tumor growth *in vivo*. Together, our findings support a critical role of PP2A in the prevention of PDAC nutrient scavenging through inhibition of the macropinosome-lysosome fusion.

## RESULTS

### PP2A promotes vesicle accumulation in PDAC cells

To determine the impact of PP2A signaling on macropinocytosis regulation, we treated a panel of human PDAC cell lines with DT-061, a direct small molecule activator of PP2A^22^. Within one-hour, high contrast vesicles were observed in DT-061 treated cells but not in the vehicle conditions in all the cell lines tested (Figure 1A). To determine if DT-061-induced vesicles are macropinosomes, PANC-1 and MIA PaCa-2 cell lines were pretreated with 5-(N-ethyl-N-isopropyl) amiloride (EIPA), a Na^+^/H^+^ exchanger inhibitor^23^, prior to the addition of DT-061. Inhibition of Na^+^/H^+^ exchange by EIPA has been shown to inhibit macropinocytosis with limited effects on other endocytic pathways^24^. Neither vehicle nor EIPA-treated cells accumulated vesicles; however, EIPA pretreatment prevented vesicle formation in DT-061 conditions, suggesting that these vesicles are macropinosomes (Figure 1B, 1C, Figure S1). To test if DT-061-mediated vesicle formation functions through PP2A, we used DT-766, a structurally similar compound to DT-061 that binds to the PP2A A/C dimer but fails to activate PP2A^25^. Consistent with an inactive compound, we did not observe vesicles in cells treated with DT-766. However, treatment with DT-766 prior to DT-061 treatment abrogated vesicle formation, suggesting that this phenotype is specific to PP2A (Figure 1D, 1E). Together, these results suggest that pharmacological PP2A activation promotes the accumulation of vesicles resembling macropinosomes in PDAC cell lines.

**Figure 1.**
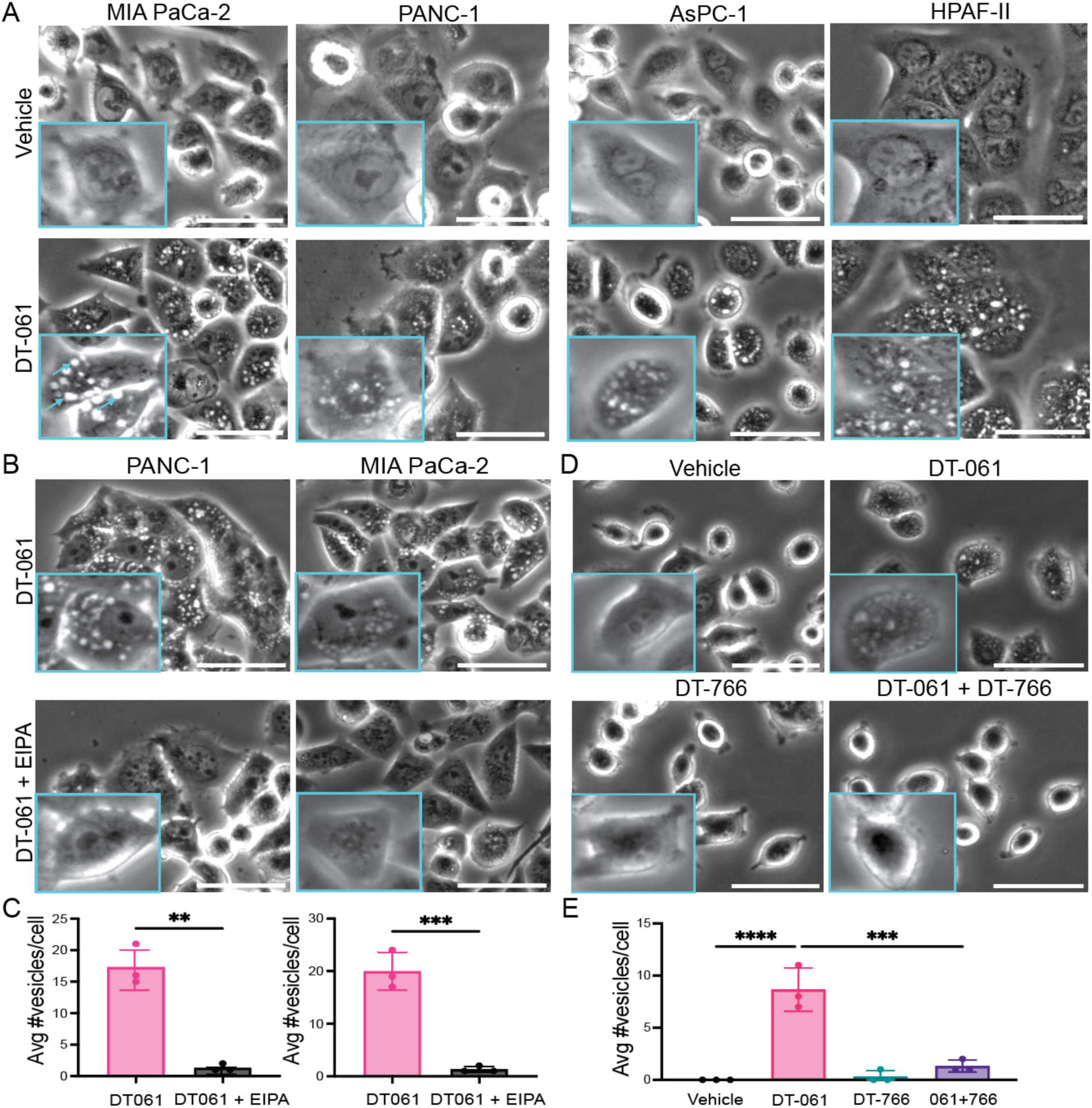
: Direct pharmacological activation of PP2A by DT-061 promotes vesicle accumulation in PDAC cells. A) Representative brightfield images of a panel of PDAC cell lines showing vesicle accumulation with 10μM DT-061 treatment for one hour compared to the vehicle control (DMSO). Arrows represent the vesicles observed in response to DT-061. B) Representative brightfield images of vesicles in PANC-1 and MIA PaCa-2 cell lines treated with 5μM DT-061 with/without 50 μM EIPA pre-treatment for 2 hours (See also Figure S1). C) Quantification of B showing average number of vesicles per cell (n=3 biological replicates) D) Representative brightfield images of AsPC-1 cells treated with single agents (vehicle control (DMSO), 20μM DT-061, 40μM DT-766) for 30 minutes or pretreated with 40μM DT-766 for 30 minutes and subsequently treated with DT-061 (combination) for 30 minutes. E) Quantification of D showing average number of vesicles per cell (n=3 biological replicates). All scale bars = 50μm. Inset denotes digital zoom. **p<0.01, ***p<0.001, ****p<0.0001 by Student’s t test or by one-way ANOVA. Data are represented as mean ± SD.

### PP2A-mediated vesicles are macropinosomes

Macropinosomes can be detected by fluorescently labeled high molecular weight (70 kDa) dextran^10,26^, which is predominantly internalized through macropinocytosis. DT-061 treated AsPc-1 and PANC-1 cells displayed a significant increase in dextran positive vesicles compared to the vehicle treated group (Figure 2A, 2B). Dextran labeled macropinosomes were also found to co-localize with the vesicles observed in DT-061 treated cells (Figure S2). As PP2A substrate specificity is dependent on the regulatory B subunits, we then sought to identify the PP2A-B subunit responsible for macropinocytosis regulation. PP2A-B56α is a known tumor suppressor^27^ and DT-061 preferentially stabilizes B56α containing PP2A heterotrimers^22^. As there is high sequence similarity within the B56 regulatory subunit family, we interrogated four of the B56 subunits, as well as B55α, which has been previously implicated in nutrient metabolism^28^. We generated AsPC-1 cells overexpressing the B56α, B56γ, B566, B56ε or B55α subunits (Figure S3) and then assessed the uptake of high molecular weight dextran. Compared to the empty vector control (EV/psin)^29^, overexpression of the B56α subunit, but not the other B subunits tested, enhanced macropinosome accumulation in AsPC-1 cells (Figure 2C, 2D). These findings support a role for the PP2A regulatory subunit, B56α, in the accumulation of macropinosomes in PDAC cells.

**Figure 2:**
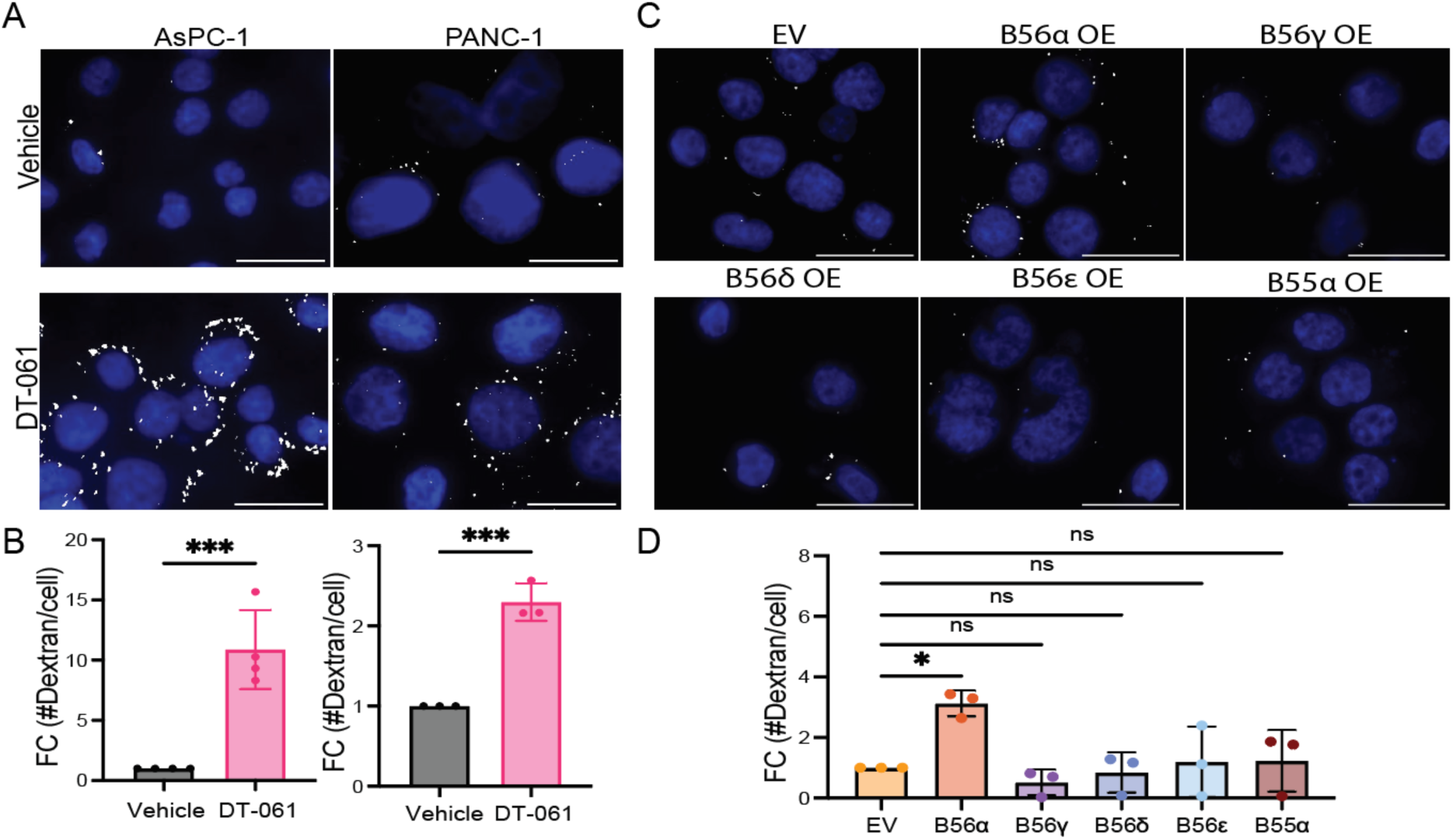
PP2A activation or overexpression of the regulatory subunit, B56α leads to macropinosome accumulation in PDAC cell lines. A) Representative fluorescence images showing Oregon Green Dextran mask (white) and DAPI (blue) in AsPC-1 and PANC-1 cells treated with control (DMSO) or 5μM DT-061 for 30 minutes. B) Quantification of A showing the fold changes in average number of Oregon green dextran puncta per cell (n=4 biological replicates for AsPC-1 and 3 biological replicates for PANC-1). C) Representative images showing Oregon Green Dextran (white) and DAPI (blue) in AsPC-1 cells overexpressing the B56 subunits (B56α, B56γ, B566 or B56ε) or B55α compared to the empty vector control (EV). D) Quantification of C showing the fold change in average number of macropinosomes per nucleus in each conditions (n= 3 biological replicates). All scale bars = 25 μm. *p<0.05, ***p<0.001 by Student’s t test or by one-way ANOVA. Data are represented as mean ± SD.

### PP2A-driven disruption of macropinocytosis maturation impairs PDAC cell viability

The Rac family small GTPases (RACs) are critical regulators of the macropinocytosis cascade^16,30^. EHT1864, a known RAC family inhibitor, has been shown to prevent the process of actin-dependent macropinocytosis^31,32^. To investigate if RAC inhibition prevents DT-061 mediated macropinosome accumulation, AsPc-1 cells were pretreated with EHT1864 and then subsequently treated with DT-061. RAC inhibition significantly prevented the DT-061-mediated increase in dextran positive macropinosomes (Figure 3A, 3B), suggesting that PP2A-mediated increase in macropinosome accumulation can be rescued by inhibiting macropinocytosis.

**Figure 3:**
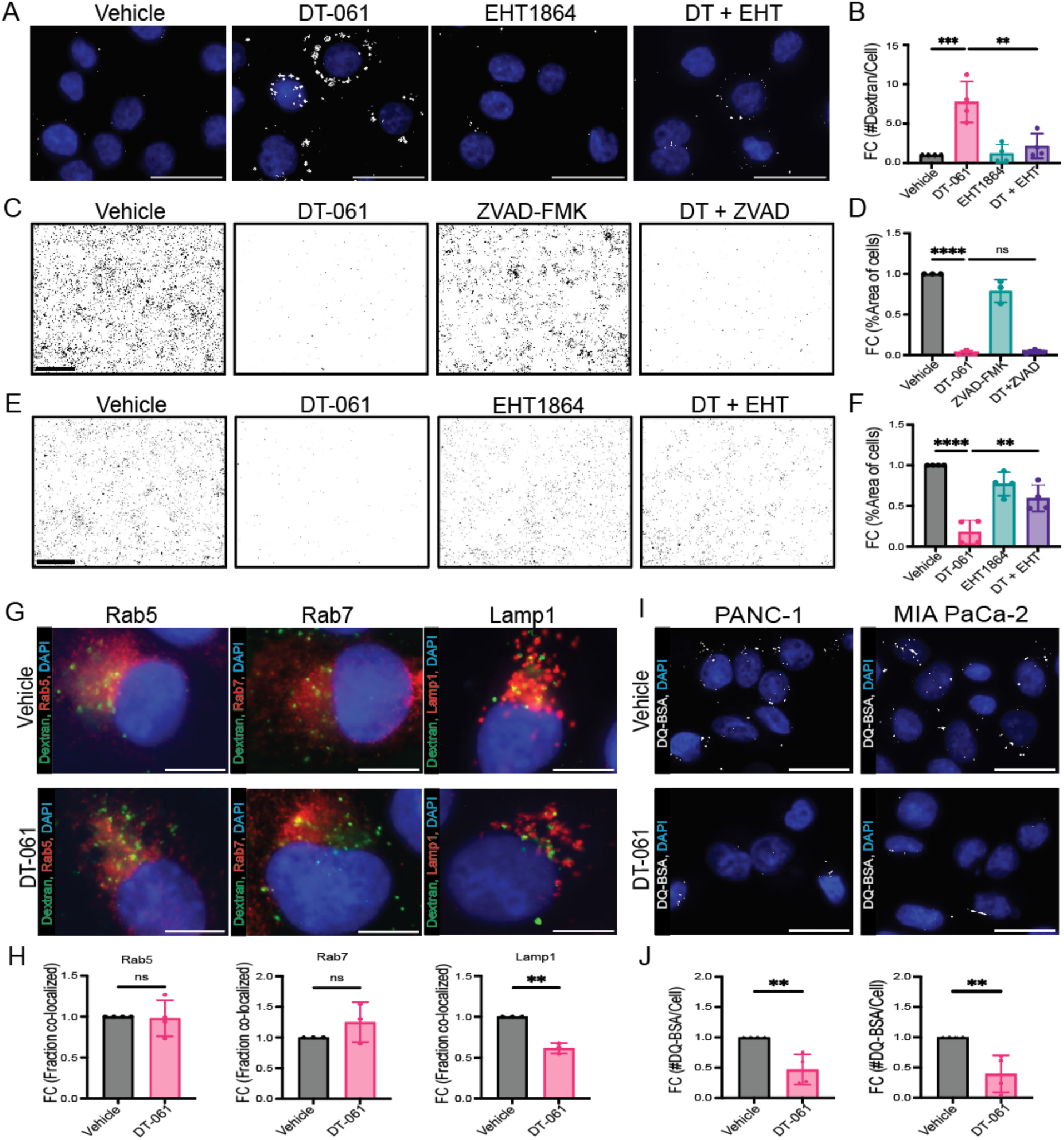
**PP2A activation decreases cell viability by preventing the fusion of macropinosomes with lysosomes**. A) Representative images showing Oregon Green Dextran (white) and DAPI (blue) in AsPC-1 cells treated with vehicle, 5μM DT-061, 25 μM EHT1864 or the combination (scale bar = 25μm). B) Quantification of A showing the fold change in average number of macropinosomes per nucleus in each conditions (n=4 biological replicates). C) Representative images of crystal violet assay in AsPC-1 cells treated with Vehicle, 5μM DT-061, 100μM ZVAD-FMK or the combination (Scale bar = 1000 μm). D) Quantification of C showing fold change in percentage occupied by stained cells (n=3 biological replicates). E) Representative images of crystal violet assay in AsPC-1 cells treated with Vehicle, 5μM DT-061, 25μM EHT-1864 or the combination (Scale bar = 1000 μm). F) Quantification of E showing fold change in percentage occupied by stained cells (n=4 biological replicates). G) Representative immunofluorescence images showing the co-localization of Oregon Green Dextran (Green), Rab5/Rab7/Lamp1 (Red) as indicated and DAPI (blue) in MIA PaCa-2 cells treated with Vehicle or 5μM DT-061 (scale bar = 10μm). H) Quantification of G showing the fold changes in fraction of co-localization of Oregon Green with the indicated markers (n=3 biological replicates). Fraction co-localized was quantified as described^21^ . I) Representative immunofluorescence images showing DQ-BSA (White) and DAPI (blue) in Vehicle or 5μM DT-061 treated PANC-1 or MIA PaCa-2 cells (scale bar = 25μm). J) Quantification of panel I showing fold change in the number of DQ-BSA puncta per cell in each condition after a chase of 1 hour (n=4 biological replicates). **p<0.01, ***p<0.001, ****p<0.0001 by Student’s t test or by one-way ANOVA. Data are represented as mean ± SD.

Macropinocytosis is known to increase PDAC cell survival^8,10^. As DT-061 increases macropinosome accumulation, we sought to determine the functional consequence of PP2A-mediated macropinosome regulation on PDAC cell survival. To assess the impact of PP2A signaling on cell viability, AsPC-1 cells were treated with vehicle or DT-061 for 24 hours and stained with crystal violet. In contrast to the known roles of macropinocytosis, DT-061 treatment significantly reduced the viability of PDAC cells compared to vehicle treated conditions (Figure 3C, 3E). Pretreatment with the pan caspase inhibitor, Z-VAD-FMK, was unable to rescue cell viability in DT-061 treated cells, potentially indicating that PP2A drives a non-programmed form of cell death (Figure 3C, 3D). Based on the increase in macropinosome accumulation, we wanted to determine the contribution of macropinocytosis to DT-061 mediated cell death. Cells were pretreated with a RAC family inhibitor, EHT1864, and then subsequently treated with DT-061. RAC inhibition significantly rescued cell viability in response to DT-061, suggesting that DT-061 mediated cell death occurs in part due to altered macropinocytosis (Figure 3E, 3F).

As dysregulation of macropinocytosis processing is known to promote a non-apoptotic form of cell death known as methuosis^33^, we investigated if the process of macropinosome maturation is impacted in response to PP2A. PDAC cells were treated with vehicle or DT-061 and immunofluorescence was performed using markers of the endocytic cascade: Rab5 (early endosomes), Rab7 (late endosomes), Lamp1 (lysosomes) and Oregon Green Dextran (macropinosome). In both vehicle and DT-061 treatment conditions, macropinosomes were found to co-localize with the early and late endosomal markers, Rab5 and Rab7. In contrast, there was a significant decrease in the co-localization of Dextran with the lysosomal marker, Lamp1, in DT-061 treated conditions (Figure 3G, 3H). As inhibition of macropinosome-lysosome fusion is known to drive methuosis^34,35^, we sought to test if this process is disrupted in cells with PP2A activation. PANC-1 and MIA PaCa2 cells were treated with vehicle or DT-061 and then incubated for 30 minutes with a high molecular weight dye-quenched BODIPY bovine serum albumin, (DQ-BSA), which is taken up through macropinocytosis and only fluoresces in the low pH of lysosomes^36^. After 30 minutes of DQ-BSA, cells were washed and incubated for 1 hour in DQ-BSA free medium and then fixed. DQ-BSA fluorescence signal was significantly reduced in DT-061 treated conditions compared to vehicle control (Figure 3I, 3J). Together, these findings indicate that PP2A activation leads to aberrant macropinocytosis through the disruption of macropinosome-lysosome fusion.

### PP2A-B56α associates with the lipid kinase, PIKfyve

PIKfyve, a phosphoinositide 5-kinase, ensures proper trafficking of cellular cargoes and is a central mediator of the endolysosomal fusion process^37^. Disruption of the activity or localization of PIKfyve is known to inhibit macropinosome-lysosome fusion^21,38^. Posttranslational regulation of PIKfyve by phosphorylation has been shown to be crucial in the regulation of its function^39^. Additionally, indirect PP2A activation has been shown to mislocalize PIKfyve^21^. However, it is unclear if PP2A directly posttranslationally regulates PIKfyve. Using the short linear motif (SLiM) discovery tool^40^, we found that PIKfyve contains multiple putative B56 binding motifs, LxxIxE^41^ near phosphorylation sites implicated in PIKfyve function^39,42,43^, suggesting that B56 subunits may interact with PIKfyve (See Table S1). To determine if PP2A-B56α interacts with PIKfyve, we performed a B56α immunoprecipitation assay in HEK293T cells transiently expressing PIKfyve-GFP. Immunoprecipitation using endogenous B56α antibody^44^ successfully pulled down PIKfyve-GFP, suggesting a direct interaction between these proteins (Figure 4A). To determine the impact of pharmacological PP2A activation on the interaction of PIKfyve and B56α, HEK293T cells were treated with vehicle or DT-061 and the interaction between B56α and PIKfyve was quantified using proximity ligation assay (PLA). DT-061 significantly increased the association of B56α with PIKfyve compared to vehicle, as seen by increased PLA puncta (Figure 4B, 4C). To identify putative phosphorylation sites on PIKfyve regulated by PP2A, HEK293T cells with transient expression of PIKfyve-GFP were treated with vehicle or DT-061 and with or without glutamine deprivation, which has been shown to promote macropinocytosis in PDAC cells^8^. PIKfyve was then immunoprecipitated with anti-GFP and subjected to mass spectrometry as described^45,46^ under these conditions. DT-061 treatment resulted in three significantly downregulated phosphoserine residues on PIKfyve (S299, S1167 and S1754) compared to vehicle treated cells (Figure 4D, 4E, Table S3), suggesting that PP2A may dephosphorylate this lipid kinase. Furthermore, out of the significantly downregulated phosphorylation sites, S299, was also downregulated under glutamine deprivation conditions, suggesting that this site may be a functionally relevant PP2A regulated phosphorylation site during macropinocytosis (Figure 4E, Table S3). Together, these findings indicate that B56α interacts with PIKfyve and that PP2A activation results in differential PIKfyve phosphorylation.

**Figure 4:**
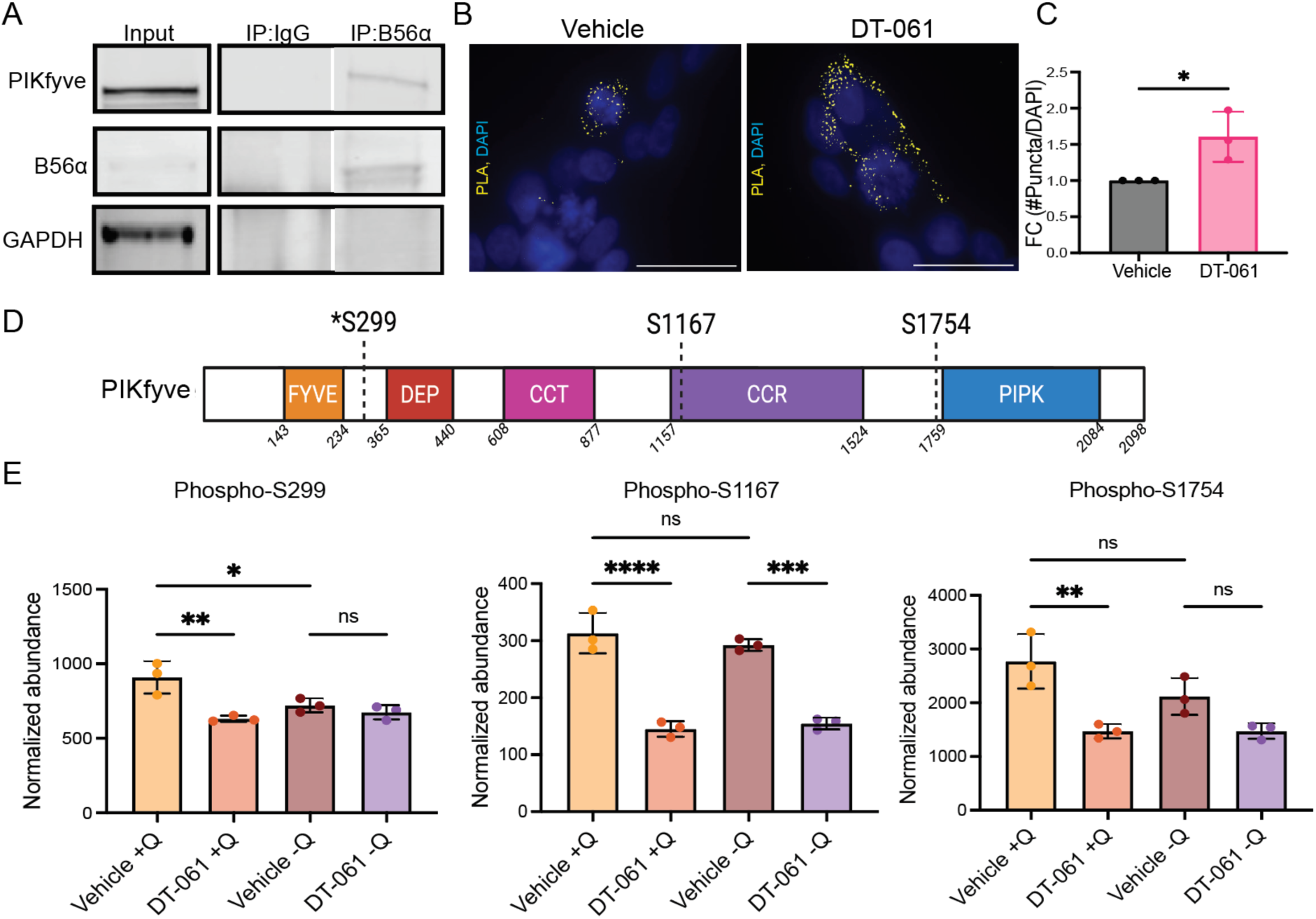
B56α interacts with PIKfyve, and the activation of PP2A downregulates PIKfyve phosphorylation on S299, S1167 and S1754. A) Immunoprecipitation blot showing the input and IP of B56α endogenous protein, where PIKfyve-GFP is pulled down with B56α in HEK293T cells expressing exogenous PIKfyve-GFP. B) Representative images of proximity ligation assay (HA & GFP) in HEK293T expressing PIKfyve-GFP and HA-B56α with Vehicle and 2.5μM DT-061 treatment for 1 hour (scale bar = 25μm). C) Quantification of B showing the fold change in number of PLA puncta per nucleus (n=3 biological replicates). D) Linear peptide map of PIKfyve showing the N-terminal FYVE domain, a Dishevelled, Egl-10 and Pleckstrin (DEP) domain, central chaperonin-like domains (CCT and CCR) and the kinase domain (PIPK). The phosphorylation sites downregulated with PP2A activation S299, S1167 and S1754 are shown in the peptide map. **\***represents the phosphorylation site with a putative B56 binding SLiM domain in close proximity. E) Quantification of the normalized abundances of differential phosphorylation sites on PIKfyve with DT-061 treatment with/without glutamine compared to vehicle control (n=3 biological replicates). *p<0.05 **p<0.01, ***p<0.001, ****p<0.0001 by Student’s t test or by one-way ANOVA. Data are represented as mean ± SD.

### PP2A activation enhances cellular responses to inhibition of nutrient metabolism

Glutamine metabolism is known to be crucial in pancreatic tumorigenesis, with low glutamine promoting macropinocytosis in PDAC^8,47^. We have determined that activation of PP2A drives aberrant macropinocytosis in PDAC cells through the disruption of macropinosome-lysosome fusion (Figures 2-4). Therefore, to identify if pharmacological activation of PP2A functions synergistically with compounds that disrupt glutamine uptake or metabolism, a panel of PDAC cells were treated with a combination of DT-061 and V-9302, CB-839, or DON^48,49^. V-9302 is an inhibitor of the amino acid transporter ASCT2 and disrupts amino acid transport into the cells^50^, CB-839 is a glutaminase (GLS) inhibitor that prevents the conversion of glutamine to glutamate^51^, and DON is a substrate analogue of glutamine^52^. Out of the three combination treatments, the majority of cell lines showed synergy with V-9302 and DT-061, as shown by high BLISS synergy and combination sensitivity scores^53^ (Figure 5A, 5B, S4). Similarly, an indirect activator of PP2A, SH-BC-893, also showed synergy with V-9302 (Figure S4). These findings indicate that the combination of PP2A activation with V-9302 effectively prevents PDAC cell survival *in vitro*.

**Figure 5:**
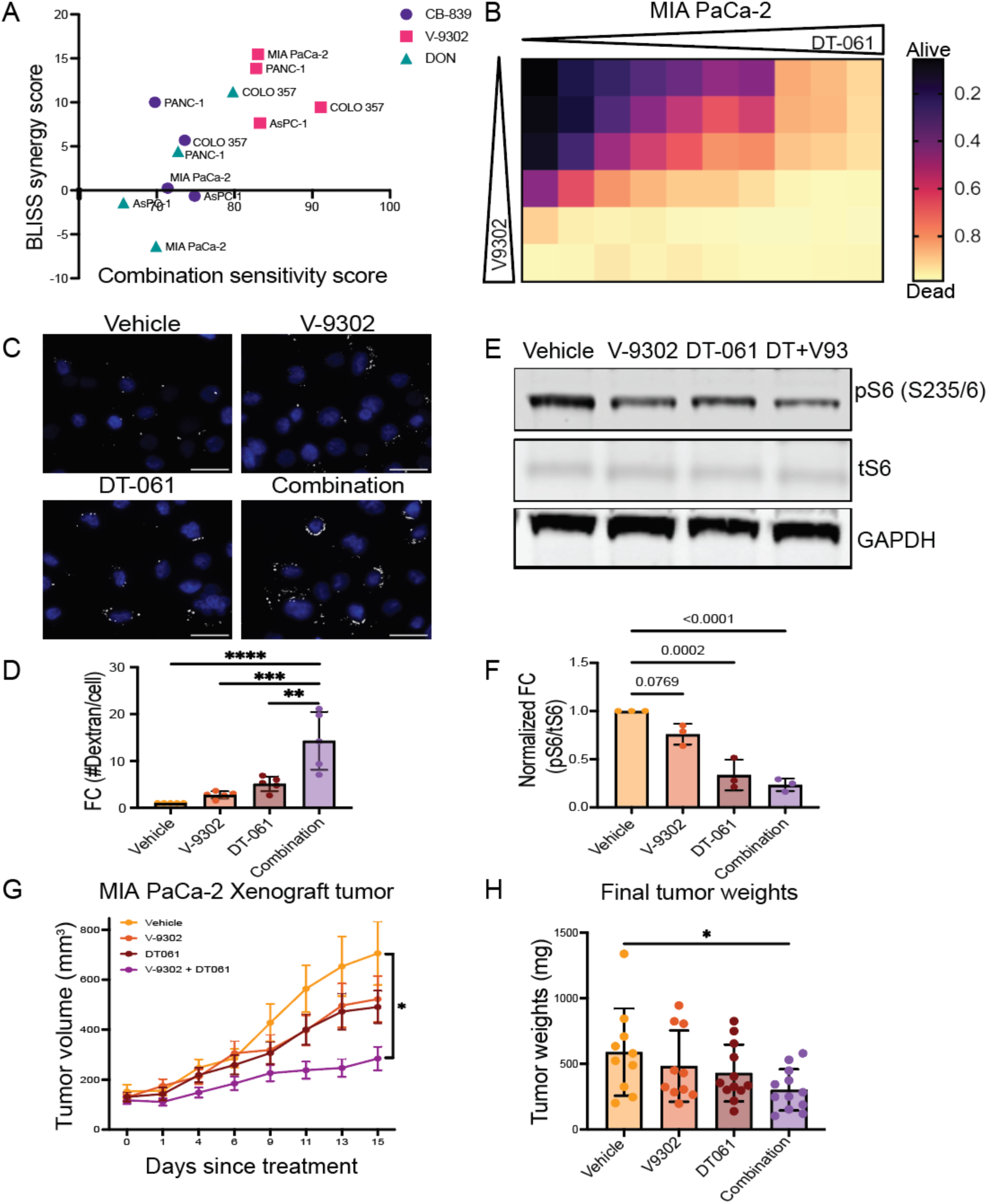
**DT-061 is synergistic with the amino acid transporter antagonist, V-9302 *in vitro* and prevents tumor progression *in vivo*** A) Plot showing the BLISS synergy and combination sensitivity scores of PDAC cell lines treated with DT-061 and glutamine inhibitors: CB-839, V-9302 or DON (n=3 biological replicates). See also Figure S4. B) Representative drug matrix showing the combination of DT-061 and V-9302 in MIA PaCa-2 cells. C) Representative images showing Oregon Green Dextran (white) and DAPI (blue) in AsPC-1 cells treated with Vehicle, 5μM DT-061, 5μM V-9302 or the combination (scale bar = 25μm). D) Quantification of C showing the fold change in average number of macropinosomes per nucleus in each condition (n=5 biological replicates). E) Representative western blot of phospho-S6, total-S6 and GAPDH proteins in AsPC-1 cell treated with Vehicle, 5μM DT-061, 2.5μM V-9302 or the combination. F) Quantification shows the fold change in protein expression of phosphorylated S6 normalized to total S6 between these conditions (n=3 biological replicates). G) Tumor growth volume over time of MIA PaCa-2 xenografts in NRG mice treated with Vehicle, DT-061, V-9302 or the combination for 15 days, showing significant attenuation of tumor progression in the combination treated tumors compared to the vehicle or single agent treatments. Variance for this graph was plotted using standard error of the mean (SEM). H) Quantification of the final tumor weights (mg) of mice in the treatment groups shown in G. *p<0.05 **p<0.01, ***p<0.001, ****p<0.0001 by Student’s t test or by one-way ANOVA. All other data are represented as mean ± SD.

To test if the combination of PP2A activation and ASCT2 inhibition increases aberrant macropinocytosis, high molecular weight dextran levels were assessed in AsPC-1 cells treated with vehicle, DT-061, V-9302, or the combination. The combination of DT-061 and V-9302 resulted in a significant increase in dextran positive vesicles compared to either vehicle or single agent treatment (Figure 5C, 5D). To test if the combination treatment of V-9302 and DT-061 impacts nutrient mediated cellular signaling, the phosphorylation of S6 kinase (pS6), a target of the serine/threonine kinase mechanistic target of rapamycin complex 1/2 (mTORC1/2)^54^, was assessed by western blot. Activation of mTORC1/2 is dictated by nutrient availability and is inhibited in response to V-9302^55^. While both V-9302 and DT-061 reduced pS6 expression, the combination treated cells showed a significant downregulation in pS6 compared to vehicle (Figure 5E, 5F). We next sought to investigate whether this treatment regimen would suppress tumor growth *in vivo*. Briefly, NOD-*Rag1*^null^ *IL2Rg*^null^ (NRG) mice were subcutaneously injected with MIA PaCa-2 cells and randomly enrolled in treatment arms when the tumors reached an average volume of ∼150mm^3^. Mice were then treated with vehicle, DT-061 (15 mg/kg, o.g. BID), V-9302 (25 mg/kg, i.p QD), or the combination for 15 days. Consistent with our *in vitro* results, combination treatment significantly reduced tumor growth over time and endpoint tumor weights compared to the vehicle control (Figure 5G, 5H). Together, these results suggest that PP2A activation is synergistic with glutamine transport inhibition in a xenograft model of PDAC.

## DISCUSSION

The dense fibrotic stroma found in PDAC tumors contributes to the harsh tumor microenvironment, leading to a depletion of key nutrients essential for tumor survival^2,5^. To circumvent this depletion, tumor cells upregulate the nutrient scavenging pathway, macropinocytosis, which increases the intracellular nutrient pool and leads to increased cell proliferation and survival^10^. Here, we have established that PP2A regulates macropinocytosis through the regulatory subunit B56α. In contrast to the known role of macropinocytosis accumulation in promoting tumorigenesis, we determined that PP2A prevents the fusion of macropinosomes with the lysosomes, leading to cell death. Overall, our studies shed light on the posttranslational signaling events that regulate macropinocytosis-mediated cell death in PDAC and implicate PP2A activators as potential macropinocytosis inhibitors.

PP2A is comprised of four regulatory subunit families with different isoforms and splice variants within each family. The expression of individual subunits, as well as their function, varies widely across cellular contexts and tissue types, highlighting the importance and complexity of the PP2A signaling network to cellular function^19,20^. Macropinocytosis is induced in response to nutrient deprivation, suggesting that this pathway functions as a key adaptation to harsh environments. We and others have shown that individual PP2A B subunits contribute to the regulation of metabolism and cell plasticity during tumor progression^28,56^. Here, we identified that B56α uniquely promotes the accumulation of macropinosomes relative to other B subunits tested. While we established that B56α plays a critical role in regulating macropinocytosis under this cellular context, we cannot rule out the contribution of other B family subunits to PDAC macropinocytosis due to dynamic changes in PP2A holoenzyme composition. Therefore, determining the direct interactors of B56α during macropinocytosis will be essential in teasing apart this critical regulation.

Aberrant regulation of macropinocytosis promotes a non-apoptotic form of cell death known as methuosis^33^. While this form of cell death is mostly known to be caused by the disruption of macropinosome-lysosome fusion, there are some studies which show that hyperactivation of macropinocytosis initiation can promote methuosis^57^. Therefore, while we demonstrate that PP2A activation disrupts macropinosome maturation, initiation of the macropinocytosis cascade could also be impacted by PP2A. In support of this hypothesis, PP2A has previously been shown to regulate actin dynamics and cell polarity under various cellular contexts^58,59^. Future studies will interrogate the impact of PP2A signaling on membrane organization and dynamic uptake of extracellular fluid.

The lipid kinase, PIKfyve is the only known source of PI(3,5)P_2_ in cells and has broadly been implicated in intracellular vesicle trafficking^37^, underscoring the importance of understanding the mechanisms regulating PIKfyve function. Recently, PIKfyve has been shown to be critical in PDAC tumorigenesis, and inhibition of this kinase in combination with KRAS inhibitors show great therapeutic response in PDAC mice models^60^. A limited number of PIKfyve phosphorylation sites (S318, S1548 and S2053) have been identified. While the function of these sites remain highly understudied, they have been implicated in cell stress and adaptive response^39,42,43^. Here we show that PIKfyve interacts with the PP2A regulatory subunit, B56α. PP2A activation was associated with a downregulation of three phosphoserine residues: S299, S1167 and S1754. PIKfyve S299 resides in close proximity to a reported B56 binding SLiM motif and the PIKfyve Dishevelled, Egl-10 and Pleckstrin (DEP) domain, which contributes to PIKfyve binding partners and localization. Additionally, S299 was also found to be downregulated under glutamine deprivation, suggesting that this is a functionally relevant phosphorylation site under nutrient stress. As there is limited literature on the function of PIKfyve posttranslational modifications, these findings raise the intriguing possibility that PIKfyve may be differentially phospho-regulated in a context specific manner. These questions highlight the need for future investigations into the functional consequence of PIKfyve S299, S1167 and S1754 phosphorylation, their specific roles in metabolic pathways, and the underlying regulatory mechanisms, which are currently under active exploration.

Glutamine is among the most significantly depleted amino acids PDAC tumors and contributes to PDAC macropinocytosis^7,8^. Here, we determined that PP2A activation functions synergistically with V-9302 across multiple PDAC cell lines. V-9302 inhibits the amino acid transporter, ASCT2, which facilitates the uptake of not only glutamine but other neutral amino acids, including alanine, serine, and cysteine^50^. As V-9302 impacts signaling beyond glutamine, these findings further support the investigation into whether PDAC cells are uniquely dependent on specific amino acids in the context of PP2A activation. As inhibition of metabolic pathways has shown limited efficacy *in vivo,* our combination studies suggest that the activation of PP2A may limit metabolic adaptation ultimately increasing the therapeutic potential of these inhibitors.

In summary, this study establishes PP2A as a critical regulator of the macropinosome processing cascade. Our findings identify PP2A mediated dephosphorylation as a critical regulator of macropinosome maturation, highlighting the need for continuous study of the posttranslational mechanisms that contributes to metabolic adaptations of cancer cells and therapeutic strategies that target this process.

## SUPPLEMENTAL INFORMATION

Supplemental Figures and Tables. Figures S1-S4 and Tables S1, S2 and S3

## ACKNOWLEDGEMENTS

We would like Dr. Egon Ogris (Max Perutz Labs, Vienna, Austria) for the generation of PP2A antibodies. We would also like to thank all the members of the BAP lab for editing of the manuscript and providing constructive feedback. This study was supported by Purdue EVPRP NIH New R01 Program (BLA-P PI, STL-N co-I), Grace M. Showalter Research Trust (BLA-P); Pancreatic Cancer Action Network (BLA-P; 22-20-ALLE), Concern Foundation (BLA-P), and American Cancer Society (BLA-P PI, STL-N co-I; RSG-24-1259215-01-MM). BNH and ID was supported by the Purdue Institute for Cancer Research (NIH grant P30 CA023168). CMP was supported by the Purdue Bilsland Dissertation Fellowship. The mass spectrometry work performed in this manuscript was done by the Indiana University School of Medicine Center for Proteome Analysis. Acquisition of the IUSM Center for Proteome Analysis instrumentation used for this project was provided by the Indiana University Precision Health Initiative. The proteomics work was supported, in part, by the Indiana Clinical and Translational Sciences Institute (Award Number UL1TR002529 from the National Institutes of Health, National Center for Advancing Translational Sciences, Clinical and Translational Sciences Award) and the P30 IU Simon Comprehensive Cancer Center Support Grant (Award Number P30CA082709 from the National Cancer Institute).

## AUTHOR CONTRIBUTION

GB, CMP, STL-N, and BLA-P designed experiments. GB, CMP, SNF, ID, BNH, LEG, JW performed and analyzed experiments. KH, EHD and ALM performed mass spectrometry sample preparation, data analysis, and methods Write-up. GN provided key resources and expertise. BLA-P secured funds and provided supervision. BLA-P and GB wrote and revised the manuscript. All authors reviewed and finalized the manuscript.

## DECLARATION OF GENERATIVE AI AND AI-ASSISTED TECHNOLOGIES IN THE WRITING PROCESS

No generative AI nor AI-assisted technologies were used in the writing process and preparation of this manuscript.

## METHODS

| REAGENT or RESOURCE | SOURCE | IDENTIFIER |
| --- | --- | --- |
| **Antibodies** |  |  |
| B56α 3A6-D3 | RAPPTA, Rasool et. al<sup>44</sup> | RRID:AB_3698067 |
| B56α 1D3-E12 | RAPPTA, Rasool et. al<sup>44</sup> | RRID: AB_3698069 |
| B55α | Cell Signaling Technology | Cat#5689S;RRID: AB_10827877 |
| GFP | Cell Signaling Technology | Cat#55494S; RRID: AB_3101977 |
| GAPDH | Thermofisher | Cat# AM4300; RRID: AB_2536381 |
| Goat anti-Rabbit IgG secondary antibody, Alexa Fluor 594 | Invitrogen | Cat# A11012; RRID: AB_3697961 |
| HA-tag | Cell Signaling Technology | Cat#3724S; RRID: AB_1549585 |
| IgG | Santa Cruz Biotechnology | Cat#sc-2025; RRID: AB_737182 |
| IRDye 680RD Goat Anti-rabbit | Licor | Cat#926-68071; RRID: AB_10956166 |
| IRDye 800CW Goat Anti-Mouse | Licor | Cat#926-32210; RRID: AB_621842 |
| Lamp1 | Cell Signaling Technology | Cat#9091S; RRID: AB_2687579 |
| PIKfyve | Cell Signaling Technology | Cat#92839S; RRID: AB_3697956 |
| Phospho-S6 | Cell Signaling Technology | Cat#5364S; RRID: AB_10694233 |
| Rab5 | Cell Signaling Technology | Cat#3547S; RRID: AB_2300649 |
| Rab7 | Cell Signaling Technology | Cat#9367S; RRID: AB_1904103 |
| S6 | Cell Signaling Technology | Cat#2317S; RRID: AB_2238583 |
| **Chemicals, peptides, and recombinant proteins** |  |  |
| Lipofectamine 3000 | Invitrogen | L3000015 |
| Polybrene | Millipore Sigma | TR1003G |
| Puromycin dihydrochloride | Fisher Bioreagents | BP2956100 |
| Dulbecco's modified Eagle Medium (DMEM) | Gibco | 11995073 |
| Opti-MEM Reduced Serum Medium | Gibco | 31-985062 |
| Matrigel | Corning | 356231 |
| Oregon Green Dextran | Invitrogen | D7173 |
| DQ-BSA Green | Invitrogen | D12050 |
| Bovine Serum Albumin (BSA) | Fisher Bioreagents | BP9703100 |
| Fetal Bovine Serum (FBS) | Fisher Scientific | FB12999102 |
| HyClone Phosphate buffered saline (PBS) | Cytiva | SH3025601 |
| 16% Paraformaldehyde solution | Electron Microscopy Sciences | 50-980-487 |
| Prolong Gold Antifade Reagent | Invitrogen | P36934 |
| DAPI | Sigma-Aldrich | D9542 |
| EHT1864 | MedChemExpress | HY-16659 |
| DT-061 | MedChemExpress | HY-112929 |
| DT-766 | Gift from Goutham Narla, Lum et. al.<sup>25</sup> | N/A |
| V-9302 | Selleckchem | S8818 |
| ZVAD-FMK | Selleckchem | S7023 |
| CB839 | Selleckchem | S7655 |
| DON | Selleckchem | S8620 |
| Kolliphor HS 15 | Sigma Aldrich | 42966 |
| Dimethylacetamide (DMA) | Fisher Chemical | D135-500 |
| Dimethyl sulfoxide (DMSO) | Fisher Bioreagents | BP231-100 |
| Pierce Protease and Phosphatase inhibitor mini tablets | Fisher Scientific | A32959 |
| XT MOPS Running Buffer | Bio-Rad | 1610788 |
| XT Reducing Agent 20X | Bio-Rad | 1610792 |
| XT Sample Buffer | Bio-Rad | 1610791 |
| XT Bis-Tris Protein Gel | Bio-Rad | 3450123 |
| Immobilon PVDF Membrane | Millipore Sigma | IPFL00010 |
| Licor TBS Blocking Buffer | Licor | NC1660550 |
| Tris (2-carboxyethyl) phosphine hydrochloride (TCEP) | Sigma Aldrich | C4706 |
| Chloracetamide (CAA) | Sigma Aldrich | C0267 |
| Trypsin/LysC Gold | Promega Corporation | V5072 |
| Trifluoroacetic acid (TFA) | Fluka | 91699 |
| Pierce C18 spin columns | ThermoFisher scientific | 89870 |
| Tandem Mass Tag Pro Reagent | ThermoFisher Scientific | 44520 |
| Waters Sep-Pak Vac cartridge | Waters | WAT054955 |
| Aurora Ultimate TS Column | Ion Opticks | AUR3-25075C18 |
| Acetonitrile | Fisher Chemical | LS122500 |
| Triton X-100 | Fisher Bioreagents | BP151-500 |
| Tween 20 | Fisher Bioreagents | BP337-500 |
| Crystal Violet | Thermo Scientific Chemicals | AC229641000 |
| Methanol | Fisher Chemical | A412P-4 |
| Acetic Acid | Fisher Chemical | A38C-212 |
| Deoxycholic acid sodium salt (DOC) | Fisher Bioreagents | BP349-100 |
| Sodium Dodecyl Sulfate (SDS) | Fisher Bioreagents | BP166-100 |
| Ethylenediaminetetraacetic acid (EDTA) | Acros Organics | 10378-23-1 |
| Tris Hydrochloride (Tris HCl) | Fisher Bioreagents | BP153-500 |
| Sodium Chloride (NaCl) | Fisher Chemical | S271-3 |
| Critical commercial assays |  |  |
| NEBuilder HiFi DNA Assembly Kit | New England Biolabs | E2621S |
| Dynabeads Co-immunoprecipitation Kit | Invitrogen | 14321D |
| Dynabeads Protein-G | Invitrogen | 10007D |
| Duolink Proximity Ligation Assay Kits | Millipore-Sigma | DUO-92008, DUO-92004 and DUO-92002 |
| DC Protein Assay Kit | Bio-Rad | 5000112 |
| Experimental models: Cell lines |  |  |
| Human: HEK293T | ATCC | CRL-3216 |
| Human: AsPC-1 | ATCC | CRL-1682 |
| Human: PANC-1 | ATCC | CRL-1469 |
| Human: MIA PaCa-2 | ATCC | CRM-CRL-1420 |
| Human: HPAFII | ATCC | CRL-1997 |
| Experimental models: Organisms/strains |  |  |
| Mouse: NOD.Cg-Rag1<sup>tm1Mom</sup> Il2rg<sup>tm1Wjl</sup>/SzJ (NRG) | Jackson Laboratory | 007799 |
| Recombinant DNA |  |  |
| V245 pCEP-4HA B56alpha | Addgene | 14532 |
| V247 pCEP 4HA B56gamma1 | Addgene | 14534 |
| V249 pCEP-4HA B56delta | Addgene | 14536 |
| V250 pCEP-4HA B56epsilon | Addgene | 14537 |
| pLCP FLAG B55 alpha | Addgene | 13804 |
| pSIN-Puro | Gift from Jason Hanna, Benton et. al<sup>29</sup> | N/A |
| PIKfyve-GFP | Addgene | 121148 |
| psPAX2 | Addgene | 12260 |
| pMD2.G | Addgene | 12259 |
| Software and algorithms |  |  |
| GraphPad Prism Version 10.3.0 | Dotmatics | N/A |
| ImageJ/Fiji Version 2.14.0/1.54p | National Institute of Health | N/A |
| Odyssey DLx Version 5.2.5 | Licor | N/A |
| Proteome Discoverer Version 2.5.0.400 | Thermo Fisher Scientific | N/A |
| Adobe Illustrator 29.6 | Adobe | N/A |
| SLiMSearch4 | Krystkowiak et. al.<sup>40</sup> | https://slim.icr.ac.uk/tools/slimsearch/input |
| SynergyFinder web application Version 3 | Ianevski et. al.<sup>53</sup> | https://synergyfinder.aittokallio.group/20250625012215037657/ |

### Experimental model and study participant details

#### Cell Lines

Human cell lines were obtained from ATCC. HEK293T cell line was initially isolated from the kidney of a female patient and contains SV40 T-antigen. AsPC-1 cell line was initially derived from ascites of a 62-year-old white, female patient, where the patient cells were injected into nude mouse and harvested after tumor formation. HPAF-II was initially isolated from the peritoneal ascitic fluid of a 44-year-old white male patient with primary pancreatic adenocarcinoma, as well as metastases to the liver, lymph nodes and diaphragm. PANC-1 cell line was initially isolated from the pancreatic duct of a 56-year-old white male with epithelioid carcinoma. MIA PaCa-2 cell line was initially derived from pancreatic tumor tissue of a 65-year-old white male.

#### Mice

Tumor xenograft study was performed in NOD.Cg-Rag1^tm1Mom^ Il2rg^tm1Wjl^/SzJ (NRG) strain of mouse, with 12 female and 20 male mice, aged 12-16 weeks. Mice were divided randomly into treatment groups, with n=8 mice (3 female, 5 male) per condition. No differences based on sex was observed (Data not shown). Animal study was performed in compliance with Purdue University guidelines after approval from Purdue Animal Care and Use Committee (IBC 19-019, Protocol 1910001962). Mice were housed in a clean room (negative for murine norovirus) with sterile conditions, 12 hour light/ 12 hour dark cycle, a temperature of 72°F and humidity of 40-60%.

#### Method Details

##### Cell Culture

AsPC-1, HPAF-II, MIA PaCa-2, PANC-1 and HEK-293T cell lines were cultured in Dulbecco’s modification of Eagle’s medium (11995073, Gibco) supplemented with 10% Fetal Bovine Serum (FB12999102, Fisher Scientific). Cell were maintained at 37°C in 5% CO_2_ condition. All cells were routinely tested for Mycoplasma using PCR-based strategy. Transient transfection of GFP-PIKfyve (#121148, Addgene) was performed in HEK-293T cells using Lipofectamine 3000 (L3000015, Invitrogen) as per manufacturers’ instructions and utilized 48 hours after transfection. Plasmids HA-B56α, HA-B56γ, HA-B566, HA-B56ε or FLAG-B55α were generated by cloning respective B subunits with HA/FLAG tags from V245 pCEP-4HA B56alpha (#14532, Addgene), V247 pCEP 4HA B56gamma1 (#14534, Addgene), V249 pCEP-4HA B56delta (#14536, Addgene), V250 pCEP-4HA B56epsilon (#14537, Addgene) or pLCP FLAG B55 alpha (#13804, Addgene) into a psin-puro vector^29^ (gift from Jason Hanna) using the NEBuilder HiFi DNA assembly kit (E2621S, New England Biolabs) following manufacturers’ protocol. Lentiviral particles were produced in HEK-293T cells transfected using Lipofectamine 3000 with 8 μg of the desired plasmids (psin/EV, HA-B56α, HA-B56γ, HA-B566, HA-B56ε or FLAG-B55α) and second-generation lentiviral packaging plasmids, 6 μg psPAX2 (#12260, Addgene) and 2 μg pMD2.G (#12259, Addgene) and filtered before use. Parental AsPC-1 cells were treated with 10 μg/ml polybrene (TR1003G, Millipore Sigma) one hour before addition of the medium containing virus particles. Following the transduction, cells were selected with puromycin (BP2956100, Fisher Bioreagents) to generate stable B56/B55 overexpressing AsPC-1 cell lines.

#### *In vitro* Oregon Green Dextran Assay

50,000 cells/well were plated on acid-washed glass cover slips in a 24 well plate with DMEM + 10% fetal bovine serum for 24 hours and subjected to serum starvation for 24 hours. After starvation, cells were incubated in serum free DMEM containing 70 kDa Dextran, Oregon Green (D7173, Invitrogen) in a final concentration of 0.5 mg/ml for 30 minutes at 37°C. Following dextran incubation, cells were washed in cold PBS four times and fixed in 4% paraformaldehyde (AA433689M, Thermo scientific Chemicals) for 13 minutes. Coverslips were counterstained with DAPI (0.5 μg/ml final concentration) (D9542, Sigma Aldrich) for 5 minutes at room temperature (RT). The cover slips were then mounted onto glass slides using prolong gold antifade reagent (P36934, Invitrogen). Cells were imaged using the Nikon Eclipse N upright fluorescence microscope or EVOS M7000 Cell Imaging System. Fluorescence images were gated in FIJI Version 2.14.0/1.54p to generate masks as previously described^26^ and the average number of macropinosomes per cell was quantified.

#### Crystal violet survival assay

50,000 cells/well AsPC-1 cells were plated in 12 well plates and subjected to serum starvation for 24 hours. Respective wells were either pre-treated with 25 μM RAC inhibitor, EHT1864 (HY-16659, MedChemExpress) or DMSO for 2 hours, or 100μM Caspase inhibitor, ZVAD-FMK (S7023, Selleckchem) or DMSO for 24 hours. Following pretreatment, 5 μM DT-061 (HY-112929, MedChemExpress) was added to the respective wells and incubated for 24 hours at 37°C. After incubation, cells were fixed and stained with 0.5% crystal violet for 1 hour at RT. Stained cells were washed twice with PBS and scanned using EVOS M7000 Cell Imaging system. The percentage occupied by cells under different treatment conditions was quantified using FIJI.

#### DQ-BSA assay

PDAC cells were plated on acid-washed glass coverslips in a 24 well plate and subjected to serum starvation for 24 hours. After starvation, cells were incubated in serum free DMEM containing DQ-BSA Green (D12050, Invitrogen) in a final concentration of 200 μg/ml. Cells were treated with either DMSO or DT-061 (5μM for MIA PaCa-2 and 10μM for PANC-1) immediately after addition of DQ-BSA and incubated for 30 minutes at 37°C. All wells were then washed with cold PBS and incubated with fresh DQ-BSA free serum free DMEM (with DMSO or DT-061) for a one-hour chase. After one hour of incubation, wells were fixed with 4% paraformaldehyde. Following fixation, all cells were counterstained with DAPI (D9542, Sigma Aldrich) for 5 minutes and mounted onto glass slides using prolong gold antifade reagent (P36934, Invitrogen). Cells were then imaged using the Nikon Eclipse N upright fluorescence microscope. Fluorescence images were gated in FIJI and average number of DQ-BSA puncta per cell was quantified.

#### Western Blot

Medium was aspirated from the cells and cells were washed twice with PBS. Cells were then harvested in buffer (20mM Tris HCl pH 7.5, 50mM NaCl, 0.5% Triton X-100, 0.5% DOC, 0.5% SDS and 1mM EDTA) supplemented with Pierce protease and phosphatase inhibitor minitablet (PIA32959, Thermo Scientific). Protein concentration was estimated using the DC protein assay kit (5000112, Bio-Rad). Protein was resuspended in a mixture of XT MOPS sample buffer and XT MOPS reducing agent and boiled at 95°C for 5 minutes. Equal amounts of protein samples were then run on an SDS-PAGE Gel (3450123, Bio-Rad) and transferred to a PVDF membrane using the Trans-Blot semi-dry transfer system (1704150, Bio-Rad). The membrane was blocked in Licor TBS blocking buffer (NC1660550, Fisher Scientific) for 1 hour at RT and incubated with primary antibodies overnight at 4°C. Membranes were incubated with housekeeping (GAPDH) antibody for 20 minutes, washed and then incubated with secondary antibodies for 1 hour at RT. The blot was then scanned using the Licor Odyssey DLx Imaging system and analyzed using the Imagestudio software version 5.2.5. Antibodies used are listed in the methods section.

#### Immunoprecipitation (IP)

IP was performed in HEK-293T cells transiently expressing exogenous GFP-PIKfyve for 48 hours. Transfected cells were lysed using the IP lysis buffer (14321D, Invitrogen) supplemented with protease and phosphatase inhibitor cocktail (PIA32959, Thermo Scientific). IgG (sc-2025, Santa Cruz Biotechnology) or N-terminal B56α 1D3-E12 (Gift from Goutham Narla) antibodies were conjugated with Dynabeads protein-G (10007D, Invitrogen) for pulldown and IP was conducted as per manufacturer’s protocol. Following the IP, beads were washed and eluted in 2.5X SDS buffer and boiled for 10 minutes at 95°C. The protein samples were then run on an SDS-gel, transferred, and processed as mentioned above.

#### Proximity Ligation Assay (PLA)

HEK-293T cells expressing exogenous GFP-PIKfyve and HA-B56α for 48 hours were treated with vehicle or 2.5μM DT-061 for 1.5 hours prior to fixation with 4%PFA for 13 minutes. Cells were then permeabilized using 0.3% Triton X solution for 20 minutes. PLA was performed using the Duolink In-Situ Red Starter Kit (DUO-92008, DUO-92004 and DUO-92002, Millipore Sigma) following manufacturers’ instructions. The antibodies used to study HA-B56α and GFP-PIKfyve interaction were, 1:100 GFP (55494S, Cell signaling technology) and 1:100 HA-tag (3724S, Cell signaling technology) diluted in antibody diluent solution. Following the amplification, cells were counterstained with DAPI (D9542, Sigma Aldrich) for 5 minutes at room temperature and mounted onto glass slides using prolong gold antifade reagent (P36934, Invitrogen). Cells were imaged using the Nikon Eclipse N upright fluorescence microscope. Fluorescence images were gated in FIJI and number of PLA puncta per cell was quantified.

#### IP-Mass Spectrometry (MS)

For identification of differential phosphorylation of PIKfyve-GFP under PP2A activation, IP was performed as described above in HEK-293T cells transiently expressing exogenous GFP-PIKfyve. Cells were cultured in DMEM with or without glutamine for 24 hours and treated with vehicle control or 10μM DT-061 for 2 hours. Following the IP with GFP antibody (55494S, Cell signaling technology), beads were washed three times in PBS, flash frozen and stored at −80°C.

#### Digestion and Peptide Purification

Sample preparation, mass spectrometry analysis, bioinformatics, and data evaluation for quantitative proteomics experiments were performed in collaboration with the Indiana University Proteomics Center for Proteome Analysis at the Indiana University School of Medicine similarly to previously published protocols^46^.

After washing, beads were covered with 8 M Urea, 100mM Tris hydrochloride, pH 8.5, reduced with 5mM tris (2-carboxyethyl) phosphine hydrochloride (TCEP, Sigma-Aldrich Cat No: C4706) for 30 minutes at room temperature to reduce the disulfide bonds. The resulting free cysteine thiols were alkylated using 10 mM choloracetamide (CAA, Sigma Aldrich Cat No: C0267) for 30 minutes at RT, protected from light. Samples were diluted to 2 M Urea with 50 mM Tris pH 8.5 and proteolytic digestion was carried out with Trypsin/LysC Gold (0.4 µg, Mass Spectrometry grade, Promega Corporation Cat No: V5072) overnight at 35 °C. After digestion, samples were quenched with 0.4% trifluoroacetic acid (TFA, v/v, Fluka Cat No: 91699).

Peptides were desalted on Pierce C18 spin columns (Cat No: 89870) with a wash 0.5% TFA followed by elution in 2 x 25 µL of 70% acetonitrile 0.1% formic acid (FA). Peptides were dried by speed vacuum and resuspended 50 mM triethylammonium bicarbonate (TEAB, from 1 M stock). Each sample was then labeled for two hours at room temperature, with 0.5 mg of Tandem Mass Tag Pro reagent (manufactures instructions Thermo Fisher Scientific, TMTpro™ Isobaric Label Reagent Set; Cat No: 44520, lot no. ZI391694, see Table S2). Samples were checked to ensure >90 % labeling efficiency and then quenched with 0.3 % hydroxylamine (v/v) at room temperature for 15 minutes. Labeled peptides were then combined and dried by speed vacuum before excess TMTpro® reagent was removed by desalting on a Waters Sep-Pak® Vac cartridge (Waters™ Cat No: WAT054955) with a 1 mL wash of water, 1 mL wash of 5% acetonitrile, 0.1% triethylamine (TEA) followed by elution for in 70% acetonitrile, with 0.1% TEA).

#### Nano-LC-MS/MS

Mass spectrometry was performed utilizing an EASY-nLC 1200 HPLC system (SCR: 014993, Thermo Fisher Scientific) coupled to Eclipse™ mass spectrometer with FAIMSpro interface (Thermo Fisher Scientific). Each multiplex was run on a 25 cm Aurora Ultimate TS column (Ion Opticks Cat No: AUR3-25075C18) in a 50 °C column oven with a 180-minute gradient. Approximately 2/120 µL was injected in technical replicate with a gradient of 8-38%B over 98 minutes; 30-95% B over 10 mins; held at 80% for 2 minutes; and dropping from 95-5% B over the final 5 min (Mobile phases A: 0.1% formic acid (FA), water; B: 0.1% FA, 80% Acetonitrile (Thermo Fisher Scientific Cat No: LS122500)). The mass spectrometer was operated in positive ion mode, default charge state of 2, advanced peak determination on, and Easy IC™ on. Three FAIMS CVs were utilized (−45 CV; −55 CV; −65CV and a technical replicate with −40 CV,−50 CV, and −60 CV) each with a cycle time of 1 s and with identical MS and MS2 parameters. Precursor scans (m/z 400-1600) were done with an orbitrap resolution of 120000, RF lens% 30, 50 ms maximum inject time, standard automatic gain control (AGC) target, minimum MS2 intensity threshold of 2.5e4, MIPS mode to peptide, including charges of 2 to 6 for fragmentation with 60 sec dynamic exclusion shared across the cycles, excluding isotopes. MS2 scans were performed with a quadrupole isolation window of 0.7 m/z, 32% HCD collision energy, 50000 resolution, 200% AGC target, dynamic maximum IT, fixed first mass of 100 m/z.

#### Mass spectrometry Data Analysis

Resulting RAW files were analyzed in Proteome Discoverer™ 2.5.0.400^45^ (Thermo Fisher Scientific) with a *Homo sapiens* UniProt reviewed proteome FASTA (downloaded 051322, 20292 sequences) plus common laboratory contaminants (73 sequences). SEQUEST HT searches were conducted with full trypsin digest, 3 maximum number missed cleavages; precursor mass tolerance of 10 ppm; and a fragment mass tolerance of 0.02 Da. Static modifications used for the search were: 1) TMTpro label on peptide N-termini, 2) TMTpro label on lysine (K), and carbamidomethylation on cysteine (C) residues. Dynamic modifications used for the search were 1) oxidation on methionine (M) residues, 2) phosphorylation on serine, threonine or tyrosine residues (S, T, Y), 3) deamidation of asparagine or arginine (Q, N), 4) acetylation on protein N-termini, 5) methionine loss on protein N-termini or 6) acetylation with methionine loss on protein N-termini. A maximum of 3 dynamic modifications were allowed per peptide. Percolator False Discovery Rate was set to a strict setting of 0.01 and a relaxed setting of 0.05. IMP-ptm-RS node was used for all modification site localization scores. Values from both unique and razor peptides were used for quantification. In the consensus workflows, peptides were normalized to the protein sequences for PIKFYVE and EGFP. Quantification methods utilized TMTpro isotopic impurity levels available from Thermo Fisher Scientific. Reporter ion quantification filters were set to an average S/N threshold of 6 and co-isolation threshold of 30%. Modified peptides were excluded from protein level quantification, normalization and roll-up. Resulting grouped abundance values for each sample type, abundance ratio values; and respective unadjusted p-values (calculated by Protein Abundance ratio with no imputation and ANOVA (individual protein)) from Proteome Discover were exported to Microsoft Excel.

#### Drug matrices

Cells were plated in 2% DMEM in 96 wells plates at 5000 cells/well and incubated for 24 hours. Cells were then treated with the indicated inhibitor, as a single agent or in combination with PP2A activator DT-061 or with DMSO control. Treatments were performed in a matrix layout where cells were treated with drugs, DT-061(8-point dilution) in combination with V-9302(5-point dilution), or CB839(5-point dilution), or DON(6-point dilution) across wells. After 72 hours, cells were stained with crystal violet and then lysed in 10% acetic acid. The absorbance was then measured at 570 nm to quantify cell viability. Synergy was calculated by Bliss synergy score using Synergyfinder^53^.

#### Xenograft tumor study

All animal studies were performed in compliance with Purdue Animal Care and Use Committee (PACUC) guidelines. MIA PaCa-2 cells were resuspended in 50/50 serum free DMEM/growth factor-reduced Matrigel (356231, Corning) and 2 million cells were injected subcutaneously injected into the flanks of NOD-*Rag1*^null^ *IL2Rg*^nul^ (NRG) mice. Tumors were calipered every other day. When the average tumor volume reached 150 mm^3^ (calculated using the formulae, volume = (length * width^2^)/2), mice were randomly enrolled into four treatment groups: Vehicle, V-9302, DT-061 and the combination of V-9302 and DT-061. 15 mg/kg DT-061 or vehicle (1:1:8 N,N dimethylacetamide (DMA): Kolliphor HS 15: PBS) was administered twice daily by oral gavage. 25 mg/kg V-9302 or vehicle (0.5:0.5:9 Dimethyl sulfoxide (DMSO): Kolliphor HS 15: PBS) was administered once daily by intraperitoneal injection. After 15 days of drug treatment, all mice were sacrificed, and the tumor tissues were harvested.

#### Immunofluorescence

50,000 MIA PaCa-2 cells were plated on glass cover slips in DMEM + 10% FBS for 24 hours. Cells were then serum starved for 24 hours and treated with vehicle or 5μM DT-061 for 30 minutes. Then, cells were incubated with Oregon Green Dextran (0.5 mg/ml) for 1 hour and fixed in 4% paraformaldehyde for 13 minutes after 4 washes in 1X PBS. After fixation, cells were washed in PBS 2 times for 5 minutes each. No detergents were used to ensure that dextran labeling is retained in the cells. Cells were then incubated in primary antibody (diluted in 3% BSA solution) overnight at 4°C. 1X PBS was used to wash the cells three times following the overnight incubation. Coverslips were then incubated with secondary antibody diluted 1:200 in 3% BSA at RT for 1 hour. Following the secondary antibody incubation, cells were washed three times with PBS for 5 minutes each. Finally, cells were counterstained with 0.5 μg/ml DAPI (D9542, Sigma Aldrich) for 5 minutes and the coverslips were mounted using Prolong gold antifade reagent (P36934, Invitrogen), and imaged on the Nikon Eclipse N upright fluorescence microscope. Antibodies used are listed in Key Resources Table. Co-localization was determined using the JACoP plugin in Fiji as described^21^.

#### Statistical considerations

Experiments were conducted in at least three independent biological replicates (n≥3) and graphs were plotted with mean ± standard deviation unless otherwise denoted in the figure legend. For conditions with two samples, statistical difference was determined using student’s t-test. Conditions with more than two samples were statistically tested using a one-way ANOVA and Tukey’s multiple comparisons test. Values less than p<0.05 were considered significant. *p<0.05 **p<0.01, ***p<0.001, ****p<0.0001. Data was analyzed using GraphPad Prism Version 10.3.0.

