## Supplemental figures and tables for "PP2A activation drives aberrant macropinocytosis and cell death in pancreatic ductal adenocarcinoma"

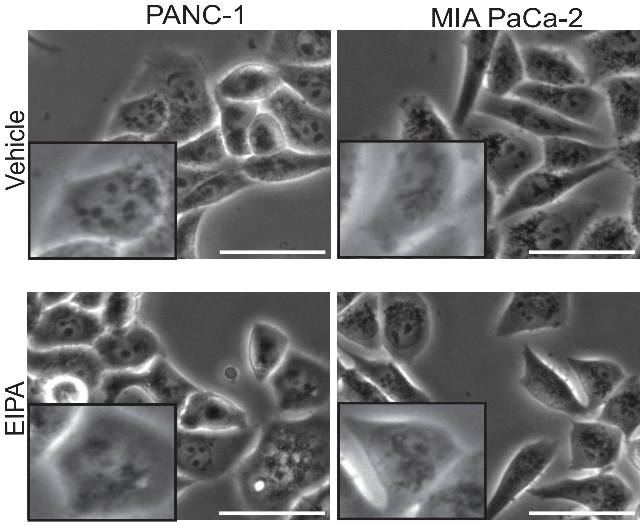


**Figure S1. Brightfield images of PANC-1 and MIA PaCa-2 cells treated with Vehicle or EIPA, related to Figure 1**

Representative brightfield images of PANC-1 and MIA PaCa-2 cells treated with vehicle (DMSO) or 50 µM EIPA. Scale bar = 50 µm. Inset denotes digital zoom.

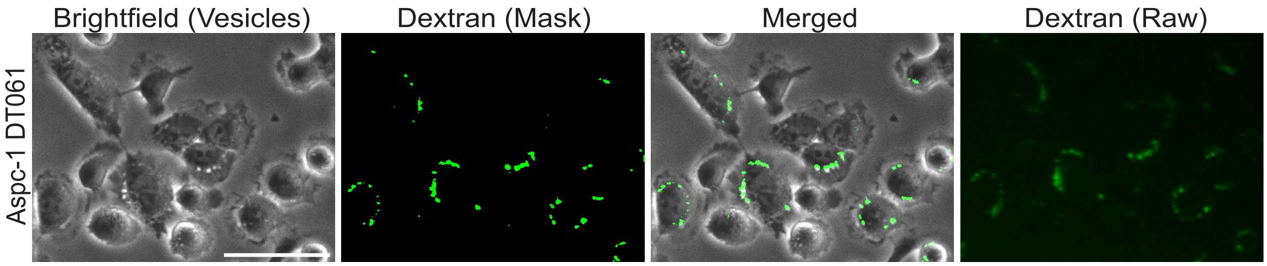


**Figure S2. Images showing the co-localization of vesicles with Dextran, related to Figure 2.**

Representative brightfield images of vesicles and Oregon Green Dextran (Raw and mask of raw images created in Fiji) in AsPC-1 cells treated with 5µM DT-061 and incubated with 0.5 mg/ml Oregon Green Dextran for 30 minutes. Images were taken post-fixation. Scale bar = 50µm.

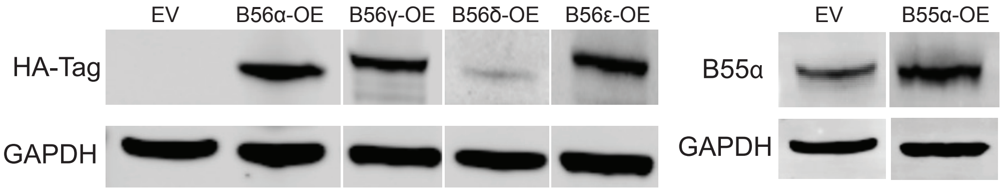


**Figure S3. Western blot showing the overexpression of B subunits, related to Figure 2.**

Representative western blot image showing the overexpression of B56 subunits, B56α, B56γ, B56δ, B56ε or B55α compared to empty vector(EV/psin), blot shows HA, B55α, and GAPDH as indicated.

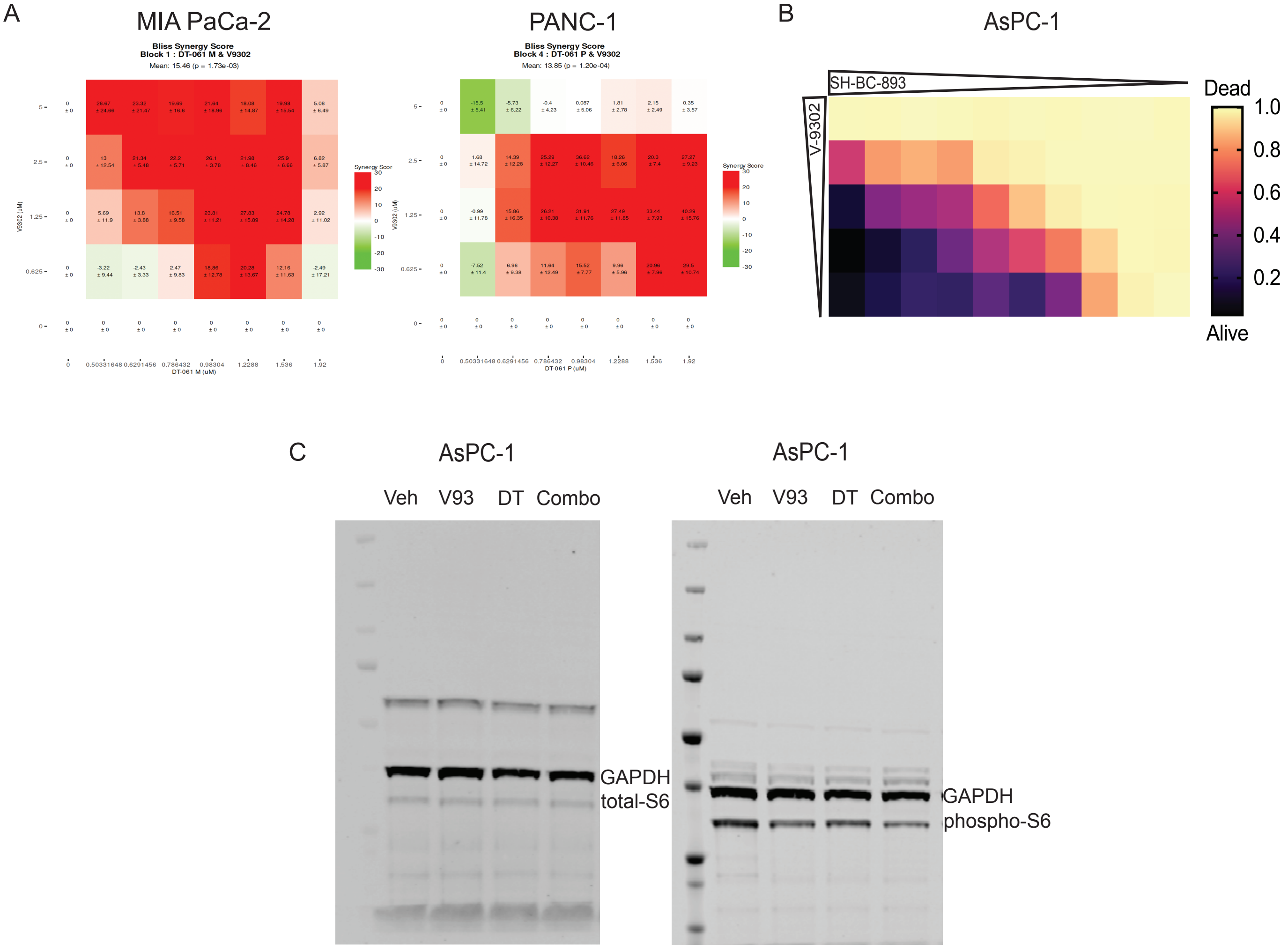


**Figure S4. Viability of AsPC-1 cells treated with increasing concentrations of V-9302 and SH-BC-893, related to Figure 5**

A) Bliss synergy scores and plot of PANC-1 and MIA PaCa-2 cells treated with a combination of V-9302 and DT-061.
B) Representative drug matrix of AsPC-1 cells treated with an indirect PP2A activator, SH-BC-893 and V-9302.
C) Representative full size blot of western blots shown in Figure 5, showing, phospho S6, total S6 and GAPDH.

| Motif Type | Motif Sequence | Start | Stop | Disorder Score |
|---|---|---|---|---|
| B56 docking motif | LKRLKEIME | 135 | 143 | 0.43 |
| B56 docking motif | LKEIMEGKS | 138 | 146 | 0.43 |
| B56 docking motif | LVSVQEDAG | 289 | 297 | 0.61 |
| B56 docking motif | LELLREENG | 592 | 600 | 0.49 |
| B56 docking motif | LARVKEILI | 849 | 857 | 0.03 |
| B56 docking motif | LRIVFEKGE | 932 | 940 | 0.38 |
| B56 docking motif | YVAIDERLA | 1424 | 1432 | 0.13 |
| B56 docking motif | LQSVFESLI | 1483 | 1491 | 0.34 |

**Table S1. Predicted B56 binding motifs on PIKfyve, related to Figure 4**

Predcited B56 binding sites on PIKfyve mined from SLiMSearch4. Eight putative B56 binding sites are found on PIKfyve. The site LVSVQEDAG from 289-297 lies in close proximity to the identified S299 phosphorylation site, which is differentially regulated with glutamine deprivation and PP2A activation (See Figure 4).

| Q status | Treatment | Replicate | TMTpro label |
|---|---|---|---|
| Plus | Vehicle | 1 | 126 |
| Plus | DT | 1 | 127N |
| Minus | Vehicle | 1 | 127C |
| Minus | DT | 1 | 128N |
| Plus | Vehicle | 2 | 128C |
| Plus | DT | 2 | 129N |
| Minus | Vehicle | 2 | 129C |
| Minus | DT | 2 | 130N |
| Plus | Vehicle | 3 | 130C |
| Plus | DT | 3 | 131N |
| Minus | Vehicle | 3 | 131C |
| Minus | DT | 3 | 132N |

**Table S2. Labeling scheme for Mass Spectrometry, related to STAR Methods**

| Confidence | Annotated Sequence | Master Protein Accessions | Master protein Descriptions | Modifications in Master Proteins | Abundance Ratio P-value (minusQ_DT) /(minusQ_Veh) | Abundance Ratio P-value (minusQ_DT) / (plusQ_DT) | Abundance Ratio P-value (minusQ_Veh) / (plusQ_Veh) | Abundance Ratio P-value (plusQ_DT) / (plusQ_Veh) |
|---|---|---|---|---|---|---|---|---|
| High | [R].IQPKNSDPFAHSK.[D] | Q9Y2I7 | 1-phosphatidylinositol 3-phosphate 5-kinase OS=Homo sapiens OX=9606 GN=PIKFYVE PE=1 SV=3 | Q9Y2I7 1xPhospho [S1167(100)] | 5.88787E-05 | 0.779375389 | 0.771786948 | 1.48991E-05 |
| High | [R].LVSVQEDAGKSPAR.[N] | Q9Y2I7 | 1-phosphatidylinositol 3-phosphate 5-kinase OS=Homo sapiens OX=9606 GN=PIKFYVE PE=1 SV=3 | Q9Y2I7 1xPhospho [S299(100)] | 0.735431228 | 0.756745595 | 0.031252533 | 0.002362718 |
| High | [R].GTAGKSPDLSSQK.[R] | Q9Y2I7 | 1-phosphatidylinositol 3-phosphate 5-kinase OS=Homo sapiens OX=9606 GN=PIKFYVE PE=1 SV=3 | Q9Y2I7 1xPhospho [S1754(100)] | 0.052245816 | 0.999997664 | 0.163375283 | 0.002415643 |
| High | [K].SQDVAVSPQQQQCSK.[S] | Q96F86 | Enhancer of mRNA-decapping protein 3 OS=Homo sapiens OX=9606 GN=EDC3 PE=1 SV=1 | Q96F86 1xPhospho [S131(100)] | 0.726033867 | 0.141200655 | 0.444784313 | 0.004166688 |
| High | [R].GTAGKSPDLSSQKR.[E] | Q9Y2I7 | 1-phosphatidylinositol 3-phosphate 5-kinase OS=Homo sapiens OX=9606 GN=PIKFYVE PE=1 SV=3 | Q9Y2I7 1xPhospho [S1754(100)] | 0.261727201 | 0.994273924 | 0.203835587 | 0.017670012 |
| High | [R].ALPNNTSSSPQPK.[K] | P04637 | Cellular tumor antigen p53 OS=Homo sapiens OX=9606 GN=TP53 PE=1 SV=4 | P04637 1xPhospho [S315(99.7)] | 0.883633346 | 0.972025059 | 0.012259774 | 0.01826 |
| High | [K].RALPNNTSSSPQPK.[K] | P04637 | Cellular tumor antigen p53 OS=Homo sapiens OX=9606 GN=TP53 PE=1 SV=4 | P04637 1xPhospho [S315(100)] | 0.299565984 | 0.969297102 | 0.006503061 | 0.045760955 |

**Table S3. Significantly altered phosphorylated peptides pulled down with GFP in PIKfyve-GFP expressing HEK-293T cells with and without DT-061 treatment in Glutamine replete (+Q) or deplete medium (-Q)**
